# confFuse: high-confidence fusion gene detection across tumor entities

**DOI:** 10.1101/163675

**Authors:** Zhiqin Huang, David T.W. Jones, Yonghe Wu, Peter Lichter, Marc Zapatka

## Abstract

**Background:** Fusion genes play an important role in the tumorigenesis of many cancers. Next-generation sequencing (NGS) technologies have been successfully applied in fusion gene detection for the last several years, and a number of NGS-based tools have been developed for identifying fusion genes during this period. Most fusion gene detection tools based on RNA-seq data report a large number of candidates (mostly false positives), making it hard to prioritize candidates for experimental validation and further analysis. Selection of reliable fusion genes for downstream analysis becomes very important in cancer research. We therefore developed confFuse, a scoring algorithm to reliably select high-confidence fusion genes which are likely to be biologically relevant.

**Results:** ConfFuse takes multiple parameters into account in order to assign each fusion candidate a confidence score, of which score ≥8 indicates high-confidence fusion gene predictions. These parameters were manually curated based on our experience and on certain structural motifs of fusion genes. Compared with alternative tools, based on 96 published RNA-seq samples from different tumor entities, our method can significantly reduce the number of fusion candidates (301 high-confidence from 8,083 total predicted fusion genes) and keep high detection accuracy (recovery rate 85.7%). Validation of 18 novel, high-confidence fusions detected in three breast tumor samples resulted in a 100% validation rate.

**Conclusions:** ConfFuse is a novel downstream filtering method that allows selection of highly reliable fusion gene candidates for further downstream analysis and experimental validations. confFuse is available at https://github.com/Zhiqin-HUANG/confFuse.

## BACKGROUND

A fusion gene is typically generated from two different genes due to genomic aberrations, or rarely at the transcript level (e.g. read-through co-transcript events). It can lead to enhanced expression or altered activity of an oncogene, or deregulation of a tumor suppressor gene [1]. Several technologies such as chromosome banding analysis and fluorescence in situ hybridization (FISH) have been successfully applied in detection of chromosomal alterations in the past (reviewed in e.g. [2]). More recently, next-generation sequencing (NGS) technologies such as paired-end RNA-seq have enabled the generation of accurate, high-resolution data in a single experiment, allowing for unbiased genome-wide fusion detection [3-7]. A great number of fusion gene detection tools/pipelines have been developed to interrogate data from NGS, particularly paired-end RNA-seq [8, 9]. The performance of the tools differs in terms of sensitivity and specificity, depending on the individual algorithms and filtering methods applied [8]. Each of these tools/pipelines has its own advantages and weaknesses. A tool/pipeline should be properly chosen for each user’s requirements, since one single tool/pipeline may not work best for all different data sets.

Fusion gene detection tools/pipelines generally consist of three major parts: firstly, mapping genomic data on reference genome/transcriptome based on existing alignment tools such as Bowtie [10, 11] and BWA [12]; second, individual methods for generating fusion candidates such as deFuse [13], FusionMap [14] and SOAPfuse [15]; and third, additional filtering algorithms to remove false positive candidates. The sensitivity of fusion gene detection mainly depends on the mapping ability in the alignment step and the specificity mostly depends on the methods of generating fusion candidates and the individual filtering methods.

Most of those tools/pipelines generate a large number of putative fusion transcripts even after filtering, of which most are likely to be false positives or of low biological interest (e.g. precursor read-through transcripts), making it hard to prioritize candidates for experimental validation. Additional filtering methods were developed based on individual datasets in order to select reliable candidates [16, 17]. Those individual filters of fusion gene candidates, however, may have a bias towards cancer or cell type-specific artifacts. A method which can work across different data sets would be very helpful for users. Some false positive fusion predictions may be due to sequencing/alignment artifacts or sequencing library preparation [18]. Furthermore, strict filtering can decrease sensitivity of true fusion detection [17]. Therefore, we developed confFuse, a new scoring algorithm, which can be applied on paired-end RNA-seq across tumor entities with both high sensitivity and high detection accuracy.

## IMPLEMENTATION

confFuse was designed to rank fusion candidates based on deFuse output by assigning each fusion candidate a confidence score, with the aim of markedly reducing the total number of fusion candidates while retaining a high recall rate for true positives. It takes multiple features into account, including some from the standard deFuse output and also newly generated features, with each given a specific score weight. The final confidence score is the sum of the score weights of different single/combined features (the initial baseline score is 10). These parameter weightings were manually optimized in comparison to a known validated fusion list, in order to achieve a balance between eliminating false positives whilst retaining true fusions. Fusion candidates scoring between 8 and 10 are considered as being high-confidence candidates. The main features used to calculate this score are described below and summarized in Suppl. Table ST1.

### Training data

Sixteen recently published pediatric glioblastoma RNA-seq samples were chosen as the first training data [19]. Fusion gene candidates in these 16 samples were first identified by tools SOAPfuse and TopHat2-Fusion [20]. High-confidence candidates were then filtered for common artifacts and by visual inspection of fusion break points of exons between two fusion partners [19]. In total, 40 fusion genes were successfully verified by RT-PCR among the 16 samples. The first training data was mainly used to select features from deFuse reports.

The second training data contains 96 RNA-seq samples from seven studies, including pilocytic astrocytoma (n=7) [21], thyroid cancer (n=5) [22], glioblastoma (n=47) [23], lung adenocarcinoma (n=28) [24], ependymoma (n=7) [25], lung cancer liver metastasis (n=1) [26], and biphenotypic sinonasal sarcoma (n=1) [27]. Those samples, including 126 experimentally verified fusion genes, were used to optimize the score weights.

### Validation data

A published study of early-onset prostate cancer including 11 RNA-seq samples were chosen for validation in silico [5] and three primary breast cancer samples were used for experimental validation.

### Artifact list

Despite a prominent role for oncogenic gene fusions in multiple cancer types, it is relatively rare for the exact same fusion to be detected across multiple, unrelated tumor entities. Fusions identified in multiple samples from different tumor entities based on currently available fusion detection tools are therefore mostly considered to be of high false positive rate. This high false positive prediction may be due to genomic complexity such as repeat regions or mapping artifacts in the alignment step. In total, 171 paired-end RNA-seq samples from 15 different entities were used to generate an artifact list of fusions identified in multiple samples from several different entities (Figure 1). To increase the sensitivity, a small number of verified fusions were manually excluded from the artifact list. We aim to assign high-confidence fusion genes a score between 8 and 10, and consider fusions contained in the artifact list to be of high false positive rate. confFuse therefore assigns a negative score (-6) to those fusions in the artifact list in order to rank them outside of the range of confident predictions.

**Figure 1:**
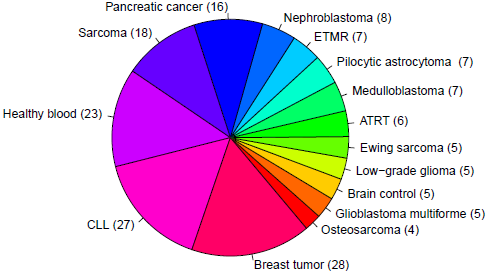
171 paired-end RNA-seq samples from 15 different entities were used to generate an artifact list of fusion genes. ETMR: embryonal tumor with multilayered rosettes; CLL: chronic lymphocytic leukemia; ATRT: atypical teratoid rhabdoid tumor.

When taking fusion candidates identified by deFuse (version v0.6.1) in no less than 3 entities (recurrence ≥ 3), 2190 fusions were included in the artifact list (Figure 2). For candidates identified in ≥ 4 and ≥ 5 entities, there are 1409 and 995 fusions in the artifact list, respectively. In this study, we chose a threshold of three entities for the final artifact list. Of note, a small number of additional putative artifacts are still identified with each increase in the number of different tumor types, suggesting that accuracy could be further improved by increasing the complexity of the data set used to generate the artifact list.

**Figure 2:**
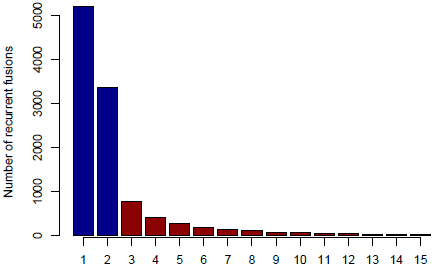
The number of recurrent fusion genes in 15 different entities. Fusions identified in more than two entities were selected for the final artifact list, resulting in 2190 artifact fusions labeled with red color.

In total, 64% (5881/9169) of putative fusion transcripts identified in the second training data were found in the artifact list. Among them, 62.3% (3666/5881) were fusions from adjacent genes and 91.5% (5378/5881) were identified by deFuse as likely being a product of alternative splicing.

### Split reads and spanning reads

One of the most important features supporting a true fusion event is the number of split reads and spanning reads. Since this is related not just to mapping performance, but also to fusion gene expression levels and sequencing depth, we found that setting a simple threshold on the number of split and spanning reads could not best distinguish true and false positive predictions. For example, a true fusion gene with low expression and low coverage sequencing depth may have only a few detectable split and spanning reads. A false positive fusion gene may have a large number of reads due to mapping artifacts and/or unreliable reads aligned to multiple genomic locations. Comparing verified fusions with all initial calls, the distribution of number of split reads and spanning reads between them is similar (Figure 3). Most of the verified fusions in the first training data have less than 200 split reads and 50 spanning reads. A threshold purely on the number of split and spanning reads therefore cannot distinguish true and false positive fusion predictions.

**Figure 3:**
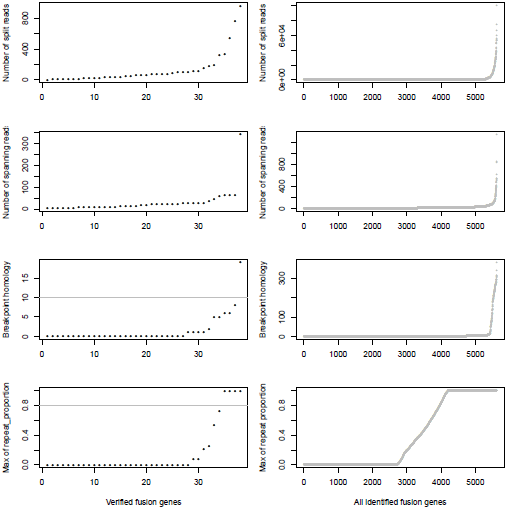
The number of split reads, spanning reads and breakpoint homology between verified and identified fusions. deFuse reported most of the verified fusion genes in 16 glioblastoma samples as containing <200 split reads, <50 spanning reads and <10 bp breakpoint homology. The maximum proportion of spanning reads in fusion partners aligned on a repeat region (repeat_proportion) is less than 80% among most of the verified fusions.

In addition, the mapping quality of reads should also be considered. Some spanning reads can be aligned to more than one genomic position, indicating reads of low mapping quality which do not reliably support a fusion event. Breakpoint homology is the number of nucleotides near the fusion break point which can map equally well to both fusion partners, with very high breakpoint homology therefore suggesting more ambiguous support for a fusion event. Most of the verified fusions contain less than 10 homologous bases at the fusion breakpoint (Figure 3). confFuse therefore assigns a negative score (-1) when breakpoint homology is ≥10. If spanning reads are mapped on a repeat region, it is difficult to identify where they are originally from. Therefore, confFuse assigns a negative score to those fusions with the majority of spanning reads aligned on repeat regions, e.g. -0.5 score for fusions of 80% up to 90% of spanning reads aligned on a repeat region (Figure 3; Suppl. Table ST1).

Fusion genes with different fusion transcripts (i.e. splice variants) in the same sample may be of high true positive rate, especially those fusion transcripts with a high count of split reads and spanning reads. We observed that deFuse sometimes reports multiple fusion transcripts for the same fusion partner genes. By combining with split reads, spanning reads and other fusion structure related features mentioned below, confFuse assigns a positive score to these fusion candidates (Suppl. Table ST1).

Taking the number of supporting reads, mapping quality and possible mapping artifacts into account, confFuse also assigns a positive score to fusions with a high number of split and uniquely mapped spanning reads or a negative score otherwise, such as -1.5 score when all the spanning reads are mapped on more than one genomic location (Suppl. Table ST1).

### Fusion structure related features

Two adjacent genes in the same orientation may give rise to an apparent fusion due to read-through transcription or aberrant splicing rather than genomic rearrangement. Although some may acquire novel function, the vast majority are expected to be false positives in terms of their biological relevance. deFuse reports an altsplice feature, indicating that a fusion may arise from alternative splicing between adjacent genes. In the first training data, verified fusion genes do not contain any read-through or alternative splicing events (Figure 4). More than 75% of initially identified fusions are, however, from an alternative splicing event. Therefore, confFuse takes those fusions with read- through or alternative splicing as high false positive fusion candidates by assigning a negative score (-4). An adjacent gene fusion ESR1:CCDC170 was reported from 22 of 990 tumor samples [28], showing the possibility of true fusions from adjacent genes. Fusion candidates from adjacent genes but without a read-through or altsplice tag are therefore given a reduced penalty of only -0.5. Furthermore, a higher ratio of inter- rather than intra-chromosomal fusions were detected in the verified fusions (Figure 4), and confFuse therefore also assigns a modest negative score (-0.5) for intrachromosomal predictions.

**Figure 4:**
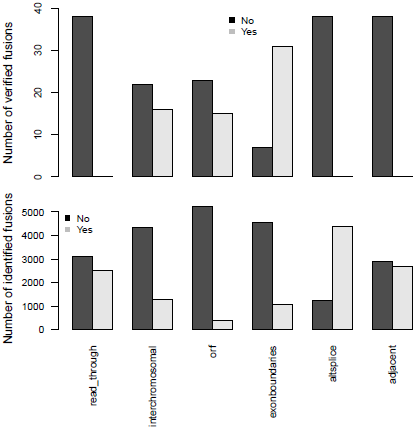
Important feature distributions between verified and identified fusions. In the first training dataset, the verified fusions (n=38) do not have any feature of read-through, alternative splicing and adjacent genes (which may also be partly due to a selection bias in those fusions selected for verification). Comparing with identified fusions, higher ratios of interchromosomal, open reading frame (orf) and exon boundaries in verified fusions were detected.

True oncogenic fusions typically preserve the open reading frame in order to form a functional fusion protein, and the precise location of a fusion breakpoint point plays a critical role in demonstrating evidence supporting true positive fusions. When the location of a fusion splicing point is at a known exon boundary, such a fusion is more likely to be a true positive. We observed higher ratios of verified fusions preserving an open reading frame and showing fusion splicing points at exon boundaries (Figure 4). confFuse assigns a negative score to fusions with non-detected open reading frame (-1 score) and with splicing point not at an exon boundary (-1.5 score). It is also more likely to be of low biological interest when a break point is located downstream of the 3’ fusion partner. confFuse takes those fusions as low-confidence ones by assigning a negative score (- 4).

## RESULTS AND DISCUSSION

### Recovery rate of verified fusions

In the first training data (n=16), 77.5% (31/40) of verified fusions were scored ≥8 by confFuse, 15% (6/40) is of 6≤ score <8, and 2.5% (1/40) is less than 6 (Suppl. Table ST2). In total, 8,083 fusion gene candidates (9,169 putative transcripts) from the second training data (n=96) were identified by deFuse using default settings, of which 126 fusions were previously validated by RT-PCR (Suppl. Table ST3). confFuse called 301 high-confidence fusion genes (score ≥ 8, 301/8,083, 3.7%). Among the 301 fusions were 108 of the 126 validated fusions, resulting in a recovery rate of 85.7% (108/126). The remaining previously validated fusions were either scored less than 8 (n=5) or were not detected or were already filtered by default deFuse parameters prior to application of confFuse (n=13) (Suppl. Table ST4). The correlation between recovery rate and confFuse score in the second training data is given in Figure 5. As annotation features such as read-through are not provided by fusionMap and soapFuse, the number of supporting reads was chosen to compare recovery rate. As the threshold for the required minimum number of supporting reads increases, the recovery rate by soapFuse and fusionMap decreases (Suppl. Figure S1). For fusions with more than five supporting reads, fusionMap predicted 964 fusions, of which 101 were previously verified, resulting in a recovery rate ~80% (101/126); soapFuse predicted 562 fusions, of which 65 were verified before, resulting in a recovery rate ~51.6% (65/126). Thus, setting a threshold only on the number of supporting reads cannot retain a high recovery rate and results in decreasing sensitivity.

**Figure 5:**
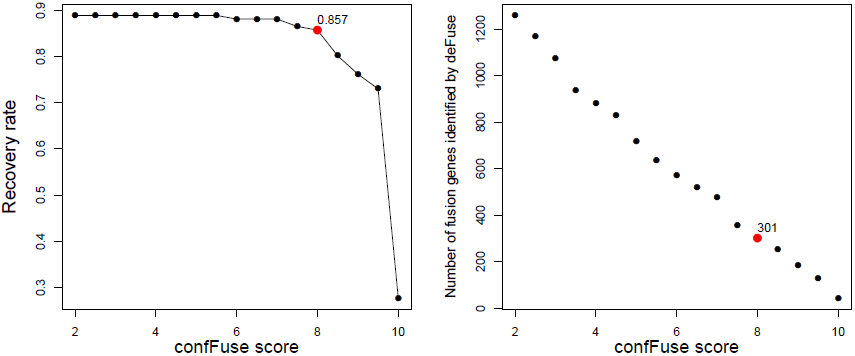
ConfFuse score and recovery rate in 96 published samples. In total, 126 fusion genes were validated, 113 of which were identified by deFuse. ConfFuse detected 108 of 126 known validated fusions with score threshold 8. The right-hand figure shows the correlation between confFuse score and the number of fusions identified by deFuse.

### Comparison of deFuse probability and confFuse confidence score

The distribution of deFuse’s own probability score was compared with our confFuse confidence score for all putative fusion transcripts in the second training data (Figure 6). Notably, there are many putative transcripts with a high deFuse probability which were assigned a low score (~ -8) by confFuse, of which most are in the artifact list or of alternative splicing feature. None of these are in the list of 126 known validated fusions in the second training data. Most of the verified fusions (108/126) are located in the range of confFuse confidence score no less than 8. It demonstrates that confFuse is able to identify the verified fusions among hundreds of putative fusions with high deFuse probability.

**Figure 6:**
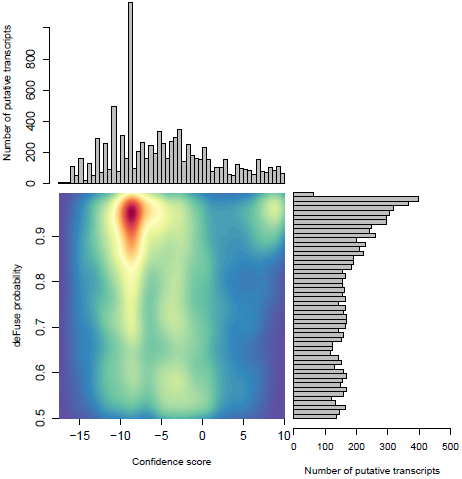
Comparison of probability predicted by deFuse and confidence score by confFuse in 96 published RNA-seq samples. High density is of red color and low density is of blue color.

### Comparison of alternative fusion detection tools

Comparing across tools, fusionMap, deFuse, deFuse-0.81 (deFuse with probability score threshold set to ≥ 0.81, as used in [13]), soapFuse, confFuse-6.5 (score ≥ 6.5) and confFuse-8 (high-confidence candidates scored ≥ 8) showed recovery rates of 91.3%, 89.7%, 84.9%, 73%, 88.1% and 85.7% respectively for the 126 validated fusions (Figure 7; Suppl. Table ST4, ST5 and ST6), indicating confFuse can dramatically reduce the number of candidates (from 8,083 to only 301) without compromising detection accuracy when compared with other available tools.

**Figure 7:**
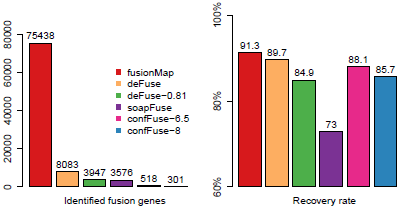
Identified fusion genes and recovery rate of validated fusions among different tools. 126 fusions were previously validated by RT-PCR. Five methods (fusionMap, deFuse, deFuse-0.81, confFuse-6.5 and confFuse-8) performed similarly in terms of recovery rate. confFuse generated much less fusion candidates than the others (higher specificity) while identifying comparable number of validated fusions (similar sensitivity).

### Validation of confFuse predicted candidates

To evaluate the accuracy of high-confidence candidate predictions (score ≥ 8), three primary breast tumor samples were sequenced to generate paired-end RNA-seq data. In total, deFuse predicted 1,026 fusion genes in the three samples, of which 18 scored ≥ 8 by confFuse. All 18 high-confidence candidates were validated with RT-PCR followed by Sanger sequencing, resulting in 100% validation rate (Figure 8; Suppl. Table ST7). To the best of our knowledge, the 18 novel fusion genes haven’t been validated before. Interestingly, one of them (QKI:PACRG) was predicted in three of 1,019 breast cancer samples from TCGA (www.tumorfusions.org), indicating that QKI:PACRG may be a novel recurrent fusion in breast cancer. QKI can suppress cell proliferation and prevent inappropriate activation of the Notch signaling pathway in lung cancer [29] and PACRG is an evolutionarily conserved protein with currently unclear function [30]. Furthermore, we randomly chose some candidates scored less than 8 for validation. Nine of 13 candidates (6.5 ≤ score ≤ 7.5, medium confidence) and three of 10 candidates (0.5 ≤ score ≤ 6, low confidence) were experimentally verified (Suppl. Table ST7). Most of the genes involved in verified fusions were in the expression level of FPKM less than 50 (median≈6.44) (Suppl. Figure SF2), indicating that it is not only very highly-expressed candidates being detected. To find out whether or not those verified fusions are individual tumor specific rather than simply artefacts of pan-breast tumor expression, the primers of verified fusions in each sample were used for validation in the other two breast primary tumor samples as control. In total, primers [31] of 14 verified fusions were used, of which 13 showed true negative in two control samples and one showed a false negative in one control sample DR8V (Suppl. Figure SF3). This ‘false negative’ fusion (CD3D:TOM1L2) in sample DR8V shows a much weaker band in gel compared with the one in sample AC72 where the fusion was true positive, and appears as a double band of different size to AC72, possibly indicating an unspecific PCR product.

**Figure 8:**
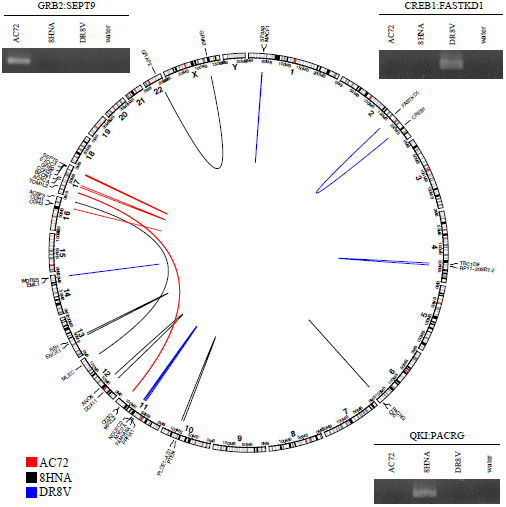
Eighteen high-confidence fusions validated by RT-PCR followed by Sanger sequencing in three primary breast tumor samples.

In addition, 11 published early-onset prostate cancer samples were used as an in silico validation dataset. 849 fusion genes were called by deFuse, 24 of which were identified as high-confidence fusion (score ≥8) by confFuse. The well-known E26 transformation-specific (ETS) fusions were detected in 10 of 11 samples, which are the same as previously published results [5]. Among the 24 fusions, 17 were confirmed by DNA-seq, FISH validation or known ETS fusions, resulting in a ~70% (17/24) recovery rate (Suppl. Table ST8).

## CONCLUSIONS

Based on deFuse reports, the scoring algorithm confFuse assigns each putative fusion transcript a confidence score. In three breast tumor samples, we achieved 100% true positive rate for high-confidence fusion candidates. Once more verified fusion genes are available as reference data, score weight optimization could be further improved. Users can also customize the score weights based on their experience to better analyze their specific data. In summary, confFuse can reliably select high-confidence fusion genes that are more likely to be biologically relevant, achieving both high validation rate and high detection accuracy, while reducing the number of candidates to a realistic number for validation.

**ABBREVIATIONS**
FPKM: Fragments Per Kilobase per Million mapped reads
Suppl: supplementary

## AVAILABILITY OF DATA AND MATERIAL

Training dataset:

Pediatric glioblastoma: EGAS00001001139 [19]

Pilocytic astrocytoma: EGAS00001000381 [21]

Thyroid cancer: SRP027364 [22]

Glioblastomas: GSE48865 [23]

Lund adenocarcinoma: ERP001058 [24]

Ependymoma: Application from the authors [25]

Lung cancer liver metastasis: ERP001058 [26]

Biphenotypic sinonasal sarcoma: GSE52257 [27]

Validation dataset:

Prostate cancer: EGAS00001000258 [5]

## AUTHORS’ CONTRIBUTIONS

ZH wrote the manuscript and implemented the method. PL, YW, DTWJ, ZH and MZ designed the experimental validation and interpreted the results. ZH designed the primers and YW performed the validation. DTWJ, YW and MZ revised the manuscript. All the authors read and approved the final manuscript.

**[H1]ACKNOWLEDGMENTS**

We would like to thank Dr. Zuguang Gu and Dr. Barbara Worst for valuable suggestions and discussion, and thank Achim Stephan for technical assistance. We also thank Prof. Dr. Roland Eils’s group for IT infrastructure support.

## CONSENT FOR PUBLICATION

Not applicable.

## ETHICS APPROVAL AND CONSENT TO PARTICIPATE

Not applicable.

## Additional Files

### Additional file 1

Table ST1: The weights of single and combined features in confFuse scoring algorithm.

Table ST2: The verified fusion genes in the first training data.

Table ST3: The verified fusion genes in the second training data.

Table ST4: The recovery of verified fusions in deFuse, confFuse-6.5 and confFuse-8.

Table ST5: The recovery of verified fusions in fusionMap.

Table ST6: The recovery of verified fusions in soapFuse.

Table ST7: The experimental validations in three breast tumor samples.

Table ST8: The high-confidence fusions in prostate samples.

Table ST9: The correlation of verified fusions and supporting reads in fusionMap.

Table ST10: The correlation of verified fusions and supporting reads in soapFuse.

### Additional file 2

Figure SF1: The correlation of recovery rate and minimum supporting reads in fusions in fusionMap and soapFuse.

Figure SF2: The expression of genes involved in verified fusions in three breast tumor samples.

Figure SF3: Validations of 14 paired primers in three breast cancer samples.

